# A new advanced backcross tomato population enables high resolution leaf QTL mapping and gene identification

**DOI:** 10.1101/040923

**Authors:** Daniel Fulop, Aashish Ranjan, Itai Ofner, Michael F. Covington, Daniel H. Chitwood, Donelly West, Yasunori Ichihashi, Lauren Headland, Daniel Zamir, Julin N. Maloof, Neelima R. Sinha

**Affiliations:** Department of Plant Biology, University of California at Davis, Davis, California; Institute of Plant Sciences and Genetics in Agriculture, The Hebrew University of Jerusalem, Rehovot, Israel

**Author notes:** Corresponding authors: Neelima R. Sinha, Department of Plant Biology, 1002 Life Sciences, One Shields Ave. Julin N. Maloof, Department of Plant Biology, 1002 Life Sciences, One Shields Ave. National Institute of Plant Genome Research, New Delhi, India (Aashish Ranjan, Neelima R. Sinha). Donald Danforth Plant Science Center, St. Louis, Missouri, United States of America. RIKEN Center for Sustainable Resource Science, Yokohama, Kanagawa, Japan (Yasunori Ichihashi). These authors contributed equally to this work (Daniel Fulop, Aashish Ranjan). **Sequence submission:** The quality filtered, barcode-sorted and trimmed short read dataset, was deposited to the NCBI Short Read Archive under accessions SRR3188298 - SRR3188738 (Bioproject accession SRP070833).

**Keywords:** heterogeneous Hidden Markov Model, regularized regression, variable selection, epistatic QTL, fine-mapping QTL

## Abstract

Quantitative Trait Locus (QTL) mapping is a powerful technique for dissecting the genetic basis of traits and species differences. Established tomato mapping populations between domesticated tomato (*Solanum lycopersicum*) and its more distant interfertile relatives typically follow a near isogenic line (NIL) design, such as the *Solanum pennellii* Introgression Line (IL) population, with a single wild introgression per line in an otherwise domesticated genetic background. Here we report on a new advanced backcross QTL mapping resource for tomato, derived from a cross between the M82 tomato cultivar and *S. pennelli*. This so-called Backcrossed Inbred Line (BIL) population is comprised of a mix of *BC*_2_ and *BC*_3_ lines, with domesticated tomato as the recurrent parent. The BIL population is complementary to the existing *S. pennellii* IL population, with which it shares parents. Using the BILs we mapped traits for leaf complexity, leaflet shape, and flowering time. We demonstrate the utility of the BILs for fine-mapping QTL, particularly QTL initially mapped in the ILs, by fine-mapping several QTL to single or few candidate genes. Moreover, we confirm the value of a backcrossed population with multiple introgressions per line, such as the BILs, for epistatic QTL mapping. Our work was further enabled by the development of our own statistical inference and visualization tools, namely a heterogeneous Hidden Markov Model for genotyping the lines, and by using state of the art sparse regression techniques for QTL mapping.

## INTRODUCTION

Leaves are the primary sites for capturing light energy, for gas exchange, and for synthesizing metabolic compounds through photosynthesis. Leaf morphology can significantly affect growth and overall performance of the plant through the interception of environmental signals (Nicotra *et al.* 2011), and is constrained by developmental, environmental, and phylogenetic contexts. Therefore, understanding the genetic and gene regulatory mechanisms underlying leaf development is of fundamental importance for increasing photosynthetic efficiency and optimizing crop performance. The basic genetic mechanisms underlying leaf development, such as leaf initiation (Waites *et al.* 1998; Timmermans *et al.* 1999; Tsiantis *et al.* 1999), establishment of leaf polarity (Kerstetter *et al.* 2001; McConnell *et al.* 2001), and leaf differentiation (Nath *et al.* 2003), have been investigated in detail in species with simple leaves (Tsukaya 2013). In contrast to the simple leaves of Antirrhinum, maize, and Arabidopsis, leaf architecture in many plants is complex with correspondingly protracted development and leaflet specification. Tomato is a classic example of a species with a complex or compound leaf. Tomato leaf lamina are subdivided into two or three orders of leaflets produced by repeated generation of auxin maxima (Koenig *et al.* 2009) that pattern leaflets along the marginal blastozone (Kim, McCormick, *et al.* 2003; Kim, Pham, *et al.* 2003), which is competent to respond to the activity of class I KNOX genes necessary for leaflet initiation (Bharathan *et al.* 2002; Efroni *et al.* 2010). Most of our current understanding of simple and compound leaf development derives from mutant analyses, expression studies, and gene expression modulation in transgenic systems (Goliber *et al.* 1999; Tsiantis and Hay 2003; Kimura *et al.* 2008). Although these studies have provided a basic understanding of leaf development, the genetic regulation of leaf development at the level of entire genomes and incorporating epistatic interactions among many different loci has only just begun to be deciphered.

Quantitative genetics, through the identification of QTL (Quantitative Trait Loci), provides an effective means to identify genetic loci regulating complex traits, such as leaf morphology, by capitalizing on existing phenotypic diversity and genetic variation. Considering the relationship of leaf traits with water use efficiency and crop yield, as well as with developmental mechanisms common to all lateral organs in plants, identification of genetic loci regulating leaf traits is of critical importance. Surprisingly, only a few systematic studies have focused on investigating the quantitative basis of leaf traits to identify genetic loci underlying yield, physiological, or morphological traits (Jiang *et al.* 2000; Pérez-Pérez *et al.* 2002; Langlade *et al.* 2005; Tian *et al.* 2011; Ku *et al.* 2012; Chitwood, Ranjan, Martinez, *et al.* 2014). Genome-wide association on a nested association mapping population identified*liguleless*, a gene regulating upright leaf angles in maize, as a factor underlying yield (Tian *et al.* 2011). However, this maize study and one recent study in *Mimulus* suggest that leaf shape is generally highly heritable and polygenic, with many additive effects, yet without much evidence for epistasis (Ferris *et al.* 2015). Nonetheless, there is a need for a comprehensive examination of the role of epistasis in regulating leaf shape and complexity in other species, particularly using statistical techniques that facilitate the inference of epistatic QTL.

Leaf traits, including leaf shape and complexity, changed repeatedly during plant evolution (Bharathan *et al.* 2002; Kim, McCormick, *et al.* 2003; Koenig and Sinha 2010; Townsley and Sinha 2012). Elucidation of the genetic basis of morphological changes that occurred during the evolution and divergence of lineages has been a major challenge in biology (Hoekstra and Coyne 2007; Carroll 2008). Phenotypic differences between populations and species can be ascribed to QTL inferred from experimental crosses, and thus analysis of QTL can address significant questions about the process of evolutionary divergence. A classic example of the use of QTL in an evolutionary context is the identification of genetic loci that are crucial to the evolution of modern maize from teosinte (Doebley and Wang 1997; Westerbergh and Doebley 2002).

The tomato clade offers an excellent system to investigate the natural variation in many plant traits, as its species have similar genetic constitution but divergent morphological and anatomical features and disparate ecologies (Stevens and Rick 1986; Moyle 2008; Ranjan *et al.* 2012). The evolution of many traits of modern domesticated tomato have been deciphered through QTL studies. Since tomato is one of the most extensively used vegetable crops in the world, most studies have focused on investigating variation in fruit size, shape and sugar content (Frary *et al.* 2000; Fridman *et al.* 2004; Xiao *et al.* 2008). Tomato is also well-established as a model system for the study of compound leaf development in Angiosperms, and the tomato clade offers a unique opportunity for investigating natural variation in leaf complexity and other leaf traits. *Solanum pennellii*, one of the most distant interfertile relatives of domesticated tomato, *Solanum lycopersicum*, possesses distinct leaf size, complexity, and morphology. In addition, *S. pennellii*shows striking differences in fruit characteristics, drought resistance, disease resistance and water use efficiency as compared to *S. lycopersicum* (Moyle 2008; Chitwood *et al.* 2013; Koenig *et al.* 2013). All aspects of leaf complexity are reduced in *S. pennellii* relative to domesticated tomato (Holtan and Hake 2003). *S. pennellii* leaves are rounder and less serrated, and leaf initiation is delayed while leaf development is accelerated relative to tomato (Ichihashi *et al.* 2014). In addition, the leaves of *S. pennellii* are more densely covered with epidermal hairs, and show a characteristic high accumulation of anthocyanin at the base of trichomes that gives the appearance of purple spots on these leaves. In spite of these dramatic differences in leaf form, only a few studies have been undertaken to identify genetic loci, and their network of downstream and epistatic interactions, that regulate the various leaf traits at the entire genome level.

As a key trait affecting crop yield, flowering time has also been investigated in tomato (Carmel-Goren *et al.* 2003; Lifschitz and Eshed 2006). The genetic control of flowering time was at first primarily elucidated in Arabidopsis, where the signals from several pathways (e.g. photoperiodic, autonomous, and vernalization, among others) are combined in a set of pathway integrator genes (*CO*, *FLC*, *SOC1*, and *FT*) that activate the floral phase transition genes *AP1* and *LFY* (Mouradov *et al.* 2002; Simpson and Dean 2002). The general manner in which these pathways are integrated to control flowering time is broadly conserved, although crop systems have been instrumental in the discovery of deviations from the Arabidopsis paradigm (Blümel *et al.* 2015). Because flowering is a major developmental phase transition, flowering time genes can strongly influence plant architecture. Tomato has played a central role in investigating how the interplay of the antagonistic roles of *SFT* and *SP* (homologs of Arabidopsis *FT* and *TFL1*, respectively) shape plant architecture in sympodial species (Lifschitz and Eshed 2006; Elitzur *et al.* 2009; Lifschitz *et al.* 2014). For instance, modulating the balance between *SFT* and *SP* significantly influences yield in tomato by controlling the number of inflorescences and their developmental timing, whereas processing tomato varieties have determinate growth as a result of being homozygous recessive sp/sp (Pnueli *et al.* 1998; Lifschitz *et al.* 2014). In contrast to its wild relatives domesticated tomato flowers rapidly and is day-neutral or photoperiod insensitive, likely as a result of artificial selection during domestication (Aung L.H. 1976; Kinet 1977; Atherton and Harris 1986; Jiménez-Gómez *et al.* 2007; Müller *et al.* 2016). For example, accession LA0716 of the wild relative *S. pennelii* flowers roughly 11 days later than tomato cultivar TM2a (deVicente and Tanksley 1993).

*Solanum lycopersicum* and *Solanum pennellii* are interfertile, enabling genetic dissection of trait differences such as those seen for leaf complexity. The interfertility of these two species was exploited to develop a genetic resource consisting of 76 introgressions lines (IL) (Eshed and Zamir 1994). Most of these introgression lines contain a single introgression from the wild species in an otherwise domesticated background. These ILs, which introgress the complete *S. pennellii*genome in the cultivated tomato (M82) background, have been extensively phenotyped for numerous traits, such as morphology, yield, fruit quality, fruit primary and secondary metabolites for identification of QTL (Rousseaux *et al.* 2005; Schauer *et al.* 2006, 2008; Semel *et al.* 2006; Stevens *et al.* 2007; Steinhauser *et al.* 2011). The striking differences in leaf morphology between *Solanum lycopersicum* and *Solanum pennellii* were capitalized on to identify QTL associated with leaf dissection (Holtan and Hake 2003). The IL population was also used to investigate the quantitative basis of adaptive evolution of leaf traits, indicating that leaf traits are an important component of climatic niche adaptation in wild tomatoes (Chitwood, Headland, Filiault, *et al.* 2012; Muir *et al.* 2014). The diverse leaf morphologies of wild tomato species (Chitwood, Headland, Kumar, *et al.* 2012) have been compared and contextualized within the shape range of the tomato IL lines and parents (Chitwood, Ranjan, Kumar, *et al.* 2014).

A few genes underlying QTL in the tomato IL population have been fine mapped and cloned. High-resolution mapping applied to the *S. pennellii* ILs led to the map-based cloning of the first two QTL: the fruit weight QTL, *fw2.2*, and the sugar yield QTL, *Brix9-2-5* (Fridman *et al.* 2000; Frary *et al.* 2000). Fine genetic mapping using the same introgression population identified the *RXopJ4* locus, a 190-kb segment on chromosome 6 of *S. pennellii*, that confers resistance to bacterial spot disease (Sharlach *et al.* 2012). Advances in next-generation sequencing and bioinformatic approaches allowed ultra high-density genotyping of the IL population, thus increasing the resolution of QTL mapping within this population (Chitwood *et al.* 2013).

Recently we used the IL population to identify more than a thousand QTL relating not only to leaf morphological features such as leaf shape, size, complexity and serration, but also to cellular features, such as epidermal cell morphology, and stomatal density and patterning (Chitwood *et al.* 2013). Though numerous QTL for various traits have been identified using the IL population, there has been limited success in fine mapping or narrowing the QTL interval to enable the identification of strong candidate genes that could then be tested for causality, in part because of the large size of most of the introgressions. Further, since almost all of the ILs contain a single introgression on a specific chromosome, the epistatic interactions regulating a trait cannot be explored using this population. Therefore, additional introgression lines, where the large introgression regions of the ILs are subdivided into smaller regions, would facilitate the identification of strong candidate genes. Moreover, lines with multiple introgressions would be greatly enable the inference of the genetic interactions regulating a trait.

Here we report on a new genetic mapping resource, an advanced backcross experimental population with ~500 Backcrossed Inbred Lines (BILs). This population was generated by repeated backcrossing (2 or 3 times) of a F1 hybrid of *S. lycopersicum*cv. M82 and *S. pennellii* and its progeny to M82, followed by selfing. For this work we sampled progeny after four or five generations of selfing (Figure 1). Ultrahigh-density genotyping using reduced representation libraries revealed that the BILs provide a much higher resolution of introgressions in terms of size compared to the ILs. In addition, most of these BILs have more than one introgression, which allows the investigation of genetic interactions underlying traits. We identified QTL regulating various leaf traits and flowering time, and also characterized potential epistatic interactions regulating these traits. Furthermore, the BIL population was also investigated for its potential to fine map the genes likely responsible for the purple spot (Punctate) phenotype of *S. pennellii*leaves, as well as to fine map leaf traits previously mapped in the ILs. Similarly, several recent studies have used a subset of the BILs with introgressions in a region of interest to further fine-map a QTL found in the ILs (Ning *et al.* 2015; Müller *et al.* 2016; Fan *et al.* 2016). The precise genotypes of these lines, their utility in fine mapping traits, the greater phenotypic spectrum seen in the lines, and their utility for mapping single locus and epistatic QTL suggests that this population will be an extremely useful resource for basic and applied research in tomato. In this study we also strove to overcome some of the limitations of QTL mapping, such as large QTL intervals encompassing many genes and methodological obstacles to revealing the genetic architecture of traits, by developing our own statistical inference and visualization tools for high precision genotyping and by using state of the art sparse regression techniques to reveal the genetic architecture of traits through the inference of epistatic interactions without preconditioning on main effect QTL.

**Figure 1.**
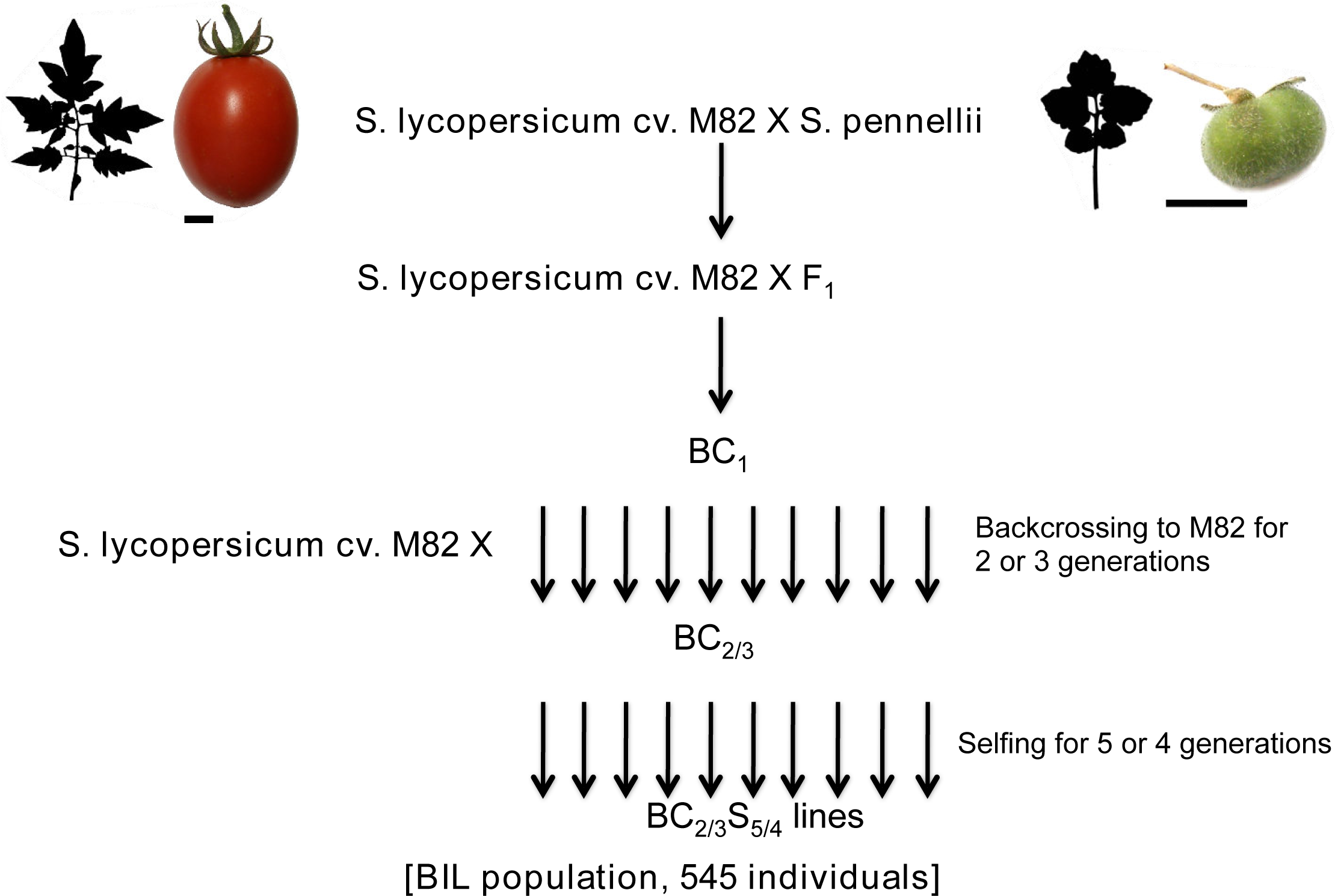
Crossing scheme for the *S. pennellii* Backcrossed Introgression Lines.

## MATERIALS AND METHODS

### Plant materials, growth conditions, experimental design, and phenotyping

Back-crossed Introgression Line (BIL) seeds were generously provided by Itai Ofner and Dani Zamir (Hebrew University, Rehovot, Israel). These BILs were generated by crossing the wild species *S. pennellii* LA0716 with the tomato cultivar M82 and backcrossing the resulting hybrid to M82 two to three times. Five and four rounds of consecutive selfing (respectively for BC_2_ and BC_3_ lines) gave rise to the BIL population assayed in this study, which consisted of approximately 545 lines.

Seeds were washed in 50% bleach for ~2 minutes, rinsed, and placed on water-soaked paper towels in Phytatrays (Sigma) to germinate. Seeds were stratified in darkness at room temperature for three days before moving to a 16:8 light cycle in growth chambers for four days. Seedlings were then transplanted into 5 x 10 subdivided trays (11’’ x 22’’ in dimension) with Sunshine Mix soil (Sun Gro) and grown in a greenhouse. Approximately three weeks after transplantation, trays were moved to a lath house where seedlings were hardened by vigorous top watering and allowing the soil to dry between waterings.

In early May 2011, hardened seedlings were transplanted into a field in Davis, CA (USA). The hypocotyl of the seedlings was buried in soil. Seedlings were initially sprinkler-watered, but later transitioned to ditch irrigation. Three randomized blocks were planted adjacent to each other, each of which contained all 545 BIL genotypes once and each of the *S. lycopersicum* cv. M82 and *S. pennellii* LA0716 parents five times.

Phenotyping for the *Punctate* locus was performed on seedlings (when the phenotype is most penetrant) at the 2-3 leaf stage when the plants were still in the greenhouse. Seedlings were scored for the punctate phenotype as a discrete, presence/absence trait. Most genotypes were scored as absent or present in all three replicates, but a few were not, and this information was noted and used during map-based cloning of the trait.

Leaf complexity was measured in the field by pairs of researchers each counting leaflets on four leaves per plant, a pair of leaves per person. Primary, intercalary, secondary, and total leaflet counts were noted. Leaflet shape was measured as previously described (Chitwood *et al.* 2013). Briefly, for each plant, the terminal and two distal lateral leaflets for five leaves per plant were placed into plastic Ziploc bags and transported back to lab. The three leaflets for each of five leaves per plant were dissected and arranged under non-reflective glass. A copy stand (Adorama, 36’’ Deluxe Copy Stand) with mounted camera (Olympus SP-500 UZ) controlled remotely using Cam2Com software (Sabsik) was used to photograph images. ImageJ (Abramoff et al., 2004) was used to isolate and threshold individual leaflets, which were saved as individual files. Leaflet outlines were then batch measured in ImageJ for area, aspect ratio, circularity, roundness and solidity values. Leaflet outlines were then processed using the software package SHAPE (Iwata and Ukai 2002) where outlines were converted to chain code, normalized Elliptical Fourier Descriptors (EFDs) calculated relative to the orientation of the proximal-distal axis of the leaflet, and a Principal Component Analysis (PCA) performed on coefficients from the first 20 harmonics.

### DNA library preparation and sequencing

After phenotyping, developing leaves and shoot apices were collected for DNA isolation and library preparation for genotyping. DNA was isolated using the DNeasy Plant Mini Kit (QIAGEN) and reduced representation libraries were prepared using restriction enzyme based RESCAN method (Monson-Miller *et al.* 2012). RESCAN library preparation involved restriction digestion of genomic DNA with the restriction enzyme NlaIII followed by adapter ligation and PCR enrichment. The PCR-enriched libraries were sequenced at the Vincent J. Coates Genomics Sequencing Laboratory at UC Berkeley on the Illumina HiSeq2000 platform (Illumina, Inc., San Diego, CA) to generate 100 bp single-end reads.

### Read processing and mapping

Reads were pre-processed, quality filtered and barcode sorted using the Fastx_toolkit as described in Chitwood et al., 2013. To remove reads originating from repeat-rich genomic regions, RESCAN sequencing reads were initially mapped to the Sol Genomic Network’s tomato repeat database using BWA (BWA parameters: -e 15 -i10 -k 1 -l 25 -n 0.05) (we created the fasta file for this from the gff3 file available at ftp://ftp.sgn.cornell.edu/genomes/Solanum_lycopersicum/annotation/ITAG2.3_release/ITAG2.3_repeats.gff3). Reads not mapped to the repeat database were extracted using bam2fastq program (http://www.hudsonalpha.org/gsl/software/bam2fastq.php). Subsequently, these repeat-filtered reads were mapped to the Heinz reference genome using the same BWA parameters. Samtools (with the ’–bq 1’ option) was used to retain the reads that mapped uniquely to the reference genome.

### Polymorphism identification

We used SNPTools (https://github.com/mfcovington/SNPtools) v0.2.4 to identify SNPs and INDELs between *Solanum lycopersicum* cv. M82 and *S. pennellii*. SNPTools is a variant-detection, genotype-scoring, and visualization tool composed of Perl modules, Perl scripts, and R scripts. The use of an earlier version of SNPTools, v0.1.5, has been described elsewhere (Devisetty *et al.* 2014).

In brief, the SNPTools polymorphism identification pipeline consists of the following steps. Putative Heinz:cv M82 and Heinz:*S. pennellii* polymorphisms were identified using a pileup-comparison-based approach. Positions where the M82 and *S. pennellii* alleles differ were retained and the Heinz-specific polymorphisms were discarded. SNPTools also has a polymorphism noise reduction step. The cv M82 and *S. pennellii* sequence alignments were genotyped at each putative polymorphism. For a true cv M82:*S. pennellii* polymorphism, the large majority of reads (>90%) should match the appropriate allele for both alignments. Putative polymorphisms that contradict this expectation were discarded.

We performed an additional, non-SNPTools, polymorphism-filtering step to remove outliers in regards to allele ratios and coverage within the BIL population. The sequence alignment files from all of the BILs were merged together and genotyped using the post-noise-reduction polymorphism database.

Polymorphisms were kept if the *S. pennellii* allele ratio was between the 25th and 75th percentile (relative to the current chromosome) and the non-parental allele ratio was less than 0.025. Since a small number of positions had excessive coverage relative the rest of the genome, we kept data only for polymorphisms with a maximum coverage of 2,000 reads.

Detailed descriptions of the polymorphism identification pipeline, including the code that was executed and genotype plots at different steps in the pipeline, are available at: https://github.com/mfcovington/bil-paper/blob/master/docs/polymorphismidentification.md.

### Heterogeneous Hidden Markov Model (HMM)

M82, heterozygous, and *S. pennellii* genotype blocks in the BILs were inferred by means of a Hidden Markov Model (HMM) implemented using the StochHMM C library plus our own C++ code for the heterogeneous transition matrix (Lott and Korf 2014). The HMM may be eventually released as standalone software. Our HMM uses a heterogeneous transition matrix in which the transition probability between states (i.e., M82, heterozygous, and *S. pennellii*) depends on the genetic distance between adjacent SNPs. With the transition matrix of our HMM we in essence modeled recombination between pairs of SNPs. Given the small number of crosses and the short distance between SNPs, we made the simplifying assumption that recombination between any two adjacent SNPs in one line of the population occurred only once (i.e., in a single generation) or not at all. We tested recombination probability equations derived from classic genetic map functions and found them to be adequate, but nevertheless unstable in a number of lines with high noise and/or low sequence coverage. Hence, we explored alternative recombination probability equations with simple parameterizations that would approximate Haldane’s and Kosambi’s equations at large genetic distances yet be more stable at short distances (i.e., less prone to switch states due local noise). The cells of the transition matrix are calculated according to the following equation for the probability of changing states:

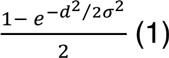
and its reciprocal for the probability of not changing states:

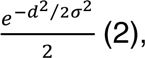
where d is genetic distance in cM between a pair of adjacent SNPs and σ is a free parameter (Figure S73). We empirically chose a value of 0.5 for σ for the present study. We plan to make the σ parameter tunable, by means of the Baum-Welch algorithm, in the training phase of the HMM in future versions of our method. Equation 1 has a low probability of transition at small genetic distances because at such distances it has an opposite curvature to Haldane’s and Kosambi’s recombination probability equations.

The prior probabilities for the hidden states are 0.88 : 0.10 : 0.02 for M82 : *S. pennellii* : heterozygous, which are roughly the expected proportion of the states in the BILs given their crossing scheme. The input SNP genotype probabilities were derived by proportionally scaling the SNP likelihoods from bcftools so that they sum to one, which implicitly assumes a uniform prior. Following previous work on genotyping HMMs (Andolfatto *et al.* 2011), an emission error rate of 1% was assumed, which captures several potential genotype error sources. The posterior decoding algorithm was used to infer posterior probabilities for each SNP sampled in a given BIL. Following convention, a posterior probability of 0.95 was used as the threshold for inferring a change in state (i.e., genotype).

In order to estimate the posterior probability of ancestry of each genomic region (i.e., which parent it came from), our HMM used the genotype probabilities at the parentally ascertained and preselected SNPs and a heterogeneous transition model, in which the transition probability between hidden states (i.e., genotypes) is a function of the genetic distance between SNPs (Figure 2). We did so in order to achieve high resolution and precision of genotype boundary calls. Common work-arounds for dealing with variable distances between SNPs within a homogeneous HMM are to bin SNPs into equidistant windows (e.g., 1 Kb in size) that are assumed to be homogenous in genotype, or to simply ignore the differences in distance and treat SNPs as if equidistant. Both of these workarounds would have degraded the resolution and precision of the inferred genotype boundaries.

**Figure 2.**
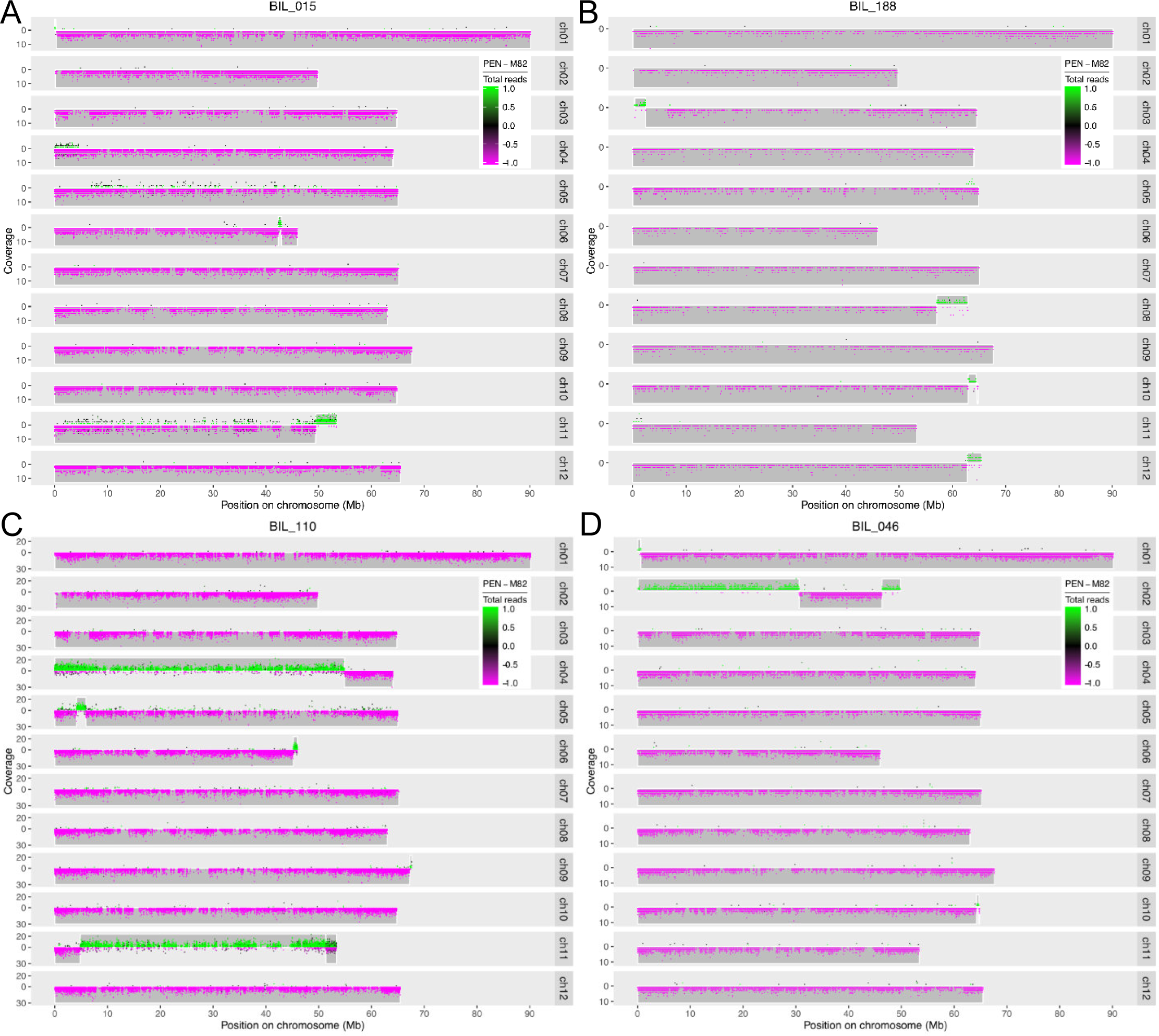
Examples of hard to genotype lines. A. High noise. B. Low sequence coverage. C. Heterozygous region with high noise. D. Narrow introgression (chromosome 10). BILs 015 and 110 were resequenced to obtain cleaner data. The boundaries obtained with the cleaner resequenced data did not differ from those shown above, demonstrating the performance of the HMM. Plots of resequenced BILs 015 and 110 are at github.com/mfcovington/bil-paper.

### BIL genotyping, plotting, and definition of bins

The BILs were genotyped using the Perl script 'extract+genotype_pileups.pl' from SNPtools v0.2.4 (https://github.com/mfcovington/SNPtools/). BIL sequence alignments were interrogated at the position of each cv M82 : *S. pennellii* polymorphism to create a set of genotype files per BIL. Recorded within a genotype file is the number of reads matching each allele for every polymorphism with read coverage on a chromosome.

To detect and fine-tune genotype bins (i.e., regions of a single parental genotype within a BIL) and the boundaries between them, we used a set of Perl scripts from detect-boundaries v0.6.1 (https://github.com/mfcovington/detect-boundaries/). The 'filter-snps.pl' script identifies bins and boundaries on a per BIL scale. The 'merge-boundaries.pl' script takes the bin and boundary data for all of the individual BILs and merges them to identify every sub-bin within the population and calculate related boundary and bin statistics. The 'fine-tune-boundaries.pl' script provides an automated approach for human verification of each boundary between bins of opposite parental genotype for a BIL. This command-line tool displays color-coded genotype data together with the currently-defined bin boundaries. Using shortcut keys, the operator can quickly and easily approve or fine-tune a boundary (at which point, the next boundary is instantly displayed for approval).

BIL genotype plots were created with the R script 'plot-genotype-and-bin.R', which uses a BIL's genotype and bin boundaries files.

### Phenotypic estimates used for mapping

BIL phenotypic estimates for plotting and QTL mapping were derived from Gaussian linear mixed effect models fitted using the lme4 R package (Bates 2015). For all traits a block random effect was used. The model used for flowering time had no additional random effects. For leaf complexity traits and asymmetric leaflet EFD principal components an additional plant level random effect was used to accommodate pseudoreplicate measurements. For shape attribute traits and symmetric leaflet EFD principal components the models included a leaflet type random effect nested within the plant level effect. For asymmetric leaflet EFD principal components, the terminal leaflet data was discarded (as these should be symmetrical or near so), and the absolute values for the left and right lateral leaflets were treated equally as they are mirror images of each other.

Model-fitted trait mean estimates were used for BIL trait plots (Figures S4-S29) and for marginal regression QTL mapping. For QTL mapping by SparseNet (see below) we generated predictions from the LMM fits such that pseudoreplication was eliminated (i.e., generated a single value per plant) and then the pseudoreplicate averaged residuals were added to the predictions.

### Heritability and Repeatability estimates

Broad sense heritability and repeatability estimates for the leaf traits were inferred by frequentist and bayesian random effect regressions using the R packages lme4 and MCMCglmm (Hadfield 2010; Bates *et al.* 2015), respectively. The experimental design random effects (e.g., block, plant, etc.) used in this case were the same as for the phenotypic estimates.

### QTL mapping

**Marginal regression:** Each individual BIL was factored for the presence or absence of *S. pennellii* introgressed DNA at a particular bin, and this process was repeated for each bin. For marginal regression, a linear model was fitted between each trait as a function of the presence for each bin. The multiple test adjustment for the significance values for each bin with respect to a given trait was performed using the Benjamini–Hochberg method. The threshold for adjusted significance value (q-value) was set at 0.001 to detect bins significantly associated with a trait.

**SparseNet:** We mapped QTL by regressing bin genotypes on phenotypic traits using the regularized regression method SparseNet (Mazumder *et al.* 2011), which uses the MC+ penalty family (an L0 to L1 family of penalties) to impose shrinkage and sparsity on the number and magnitude of regression coefficients. For each trait we performed two QTL mapping analyses with SparseNet: A) one in which we only considered non-epistatic or monogenic (single bin) QTL (which we denote as the Additive Model), B) and a second analysis in which we considered single locus QTL and all potential epistatic two-locus QTL whose bin combination were observed in the BILs (which we denote as the Epistatic Model). The Additive Model had 1,049 possible effects, i.e., equal to the number of BINs. The Epistatic Model had 307,340 possible effects, 1,049 single loci plus the 306,291 two-locus combinations observed in the population; out of the 549,676 possible two BIN combinations among the 1,049 BINs, 243,385 were not observed in the BILs.

By using a penalized method such as SparseNet we were able to map each trait on all bins and all bins plus all observed two bin combinations simultaneously. In order to choose the tuning parameter values for SparseNet, for each trait we averaged the alpha and gamma values from ten 6-fold cross-validations; the averaging was done to achieve reproducible tuning of the model. Briefly, alpha controls the severity of the penalty, and gamma controls the mixture between L0 and L1 penalties. Following cross-validation the SparseNet model was fit to all the data of a given trait using the chosen values for alpha and gamma. Because methods such as LASSO and SparseNet select a single coefficient among a set of highly correlated coefficients, we took the set of BINs with a correlation of 0.90 to a SparseNet selected BIN as a sort of empirical QTL interval within which to search for candidate genes.

### PROVEAN analysis

PROVEAN (PRotein Variation Effect ANalyzer) is a command-line tool to predict the functional impact an amino acid substitution or indel has on a protein (Choi *et al.* 2012). PROVEAN-Assembly-Line v0.3.2 (https://github.com/mfcovington/PROVEAN-Assembly-Line) was used to run a set of Perl scripts to automate high-throughput bulk PROVEAN analysis.

The 'find-aa-substitutions.pl' script was used to generate the amino acid substitution files required to run PROVEAN. The input files for this script are SNP files, a genomic FASTA reference file (S_lycopersicum_chromosomes.2.40.fa), and the corresponding GFF3 annotation file (ITAG2.3_gene_models.gff3).

The 'run-provean.pl' script was used to run the PROVEAN tool in parallel for every missense mutation, every nonsense mutation, and every mutation that caused the loss of a STOP codon. The input files for this script are the amino acid substitution files generated in the previous step and a CDS FASTA reference file (ITAG2.3_cds_SHORTNAMES.fasta).

The 'filter-provean-results.pl' was used to filter PROVEAN results for a single gene, a file containing a list of genes, or all genes within a genomic range.

### Candidate gene Identification

A set of literature-curated leaf developmental genes was used as a reference for the finding candidate genes within QTL bins for different leaf traits (Ichihashi *et al.* 2014). We selected only those genes from the set that were either differentially expressed between the two parents based on expression-QTL analysis of the IL population or that showed amino acid changes between the two parents that were predicted to be functionally significant by PROVEAN. The QTL bins were then scanned for the presence of these genes. Following, we performed permutation analyses in the R statistical environment to check whether bins associated with leaf traits were enriched for the selected literature-curated leaf developmental genes. The data were permuted 10,000 times, and the results were considered significant if < 5% of the permuted datasets equaled or exceeded the real data set.

### IL eQTL enrichment among BIL epistatic QTL

We assessed whether IL *trans*-eQTL were enriched among the set of paired regions that make up the BIL epistatic QTL using two eQTL clusters. One cluster was enriched for leaf development GO terms and the other for photosynthesis GO terms. Together these two clusters contained the majority of the literature-curated leaf development genes used for the candidate gene search (Ichihashi *et al.* 2014; Ranjan *et al.* 2016). The IRanges R/Bioconductor package (Lawrence *et al.* 2013) was used to find the overlaps between the IL *trans*-eQTL and the BIL epistatic QTL. We tested for IL *trans*-eQTL enrichment among BIL epistatic QTL by permutation using the leaf development cluster (set 1), as well as this cluster plus the photosynthesis cluster (set 2). The data were permuted 10,000 times, and the results were considered significant if < 5% of the permuted datasets equaled or exceeded the real data set.

### QTL confirmation and fine-mapping

The IRanges R/Bioconductor package (Lawrence *et al.* 2013) was used to find overlaps between the IL phenotypic QTL and the BIL single locus QTL, and the results were tabulated and plotted.

## RESULTS AND DISCUSSION

### High precision genotyping by means of a novel HMM

The Backcrossed Inbred Lines (BILs) were genotyped at high resolution using Restriction Enzyme Sequence Comparative ANalysis (RESCAN), a reduced representation genomic sequencing method (Monson-Miller *et al.* 2012; Seymour *et al.* 2012). A polymorphism database was created for sequence variants that differentiate the two parents, the domesticated tomato variety *S. lycopersicum* cv. M82 and its wild relative, *S. pennellii* LA0716. The database consisted of 346,645 polymorphisms between M82 and *S. pennellii* (295,571 SNPs and 51,074 inserted or deleted nucleotides), which was used to identify parental SNPs and INDELs for each BIL across the entire genome.

Because of the generally low and sparse sequence coverage of RESCAN, we used a novel heterogeneous Hidden Markov Model (HMM) to define the introgressed regions for each BIL. The HMM proved indispensable for accurately genotyping about 15% of the lines, which were problematic due to a variety of issues (Figure 2). Moreover, the HMM also enabled us to infer sharper recombination boundaries (github.com/mfcovington/bil-paper). The HMM inferred narrow transition boundaries in spite of noise and/or low coverage in some of the lines (Figure 2).

We were able to establish the genotype of 493 out of a total 545 BILs with confidence. However, 52 lines were found to be genotypically identical to the M82 parent, and two out of the remaining 441 were identical to two other lines. This could be due to seed or pollen contamination, and/or random effects during backcrossing and selfing. We were not able to establish the genotype of 52 other lines either due to lack of germination or excessive noise in the raw genotype data. The 439 uniquely genotyped BILs covered the entire tomato genome except for a small gap of ~400Kb on chromosome 1. We detected heterozygous introgressions on almost all chromosomes (Figures 2C, 3A, and 4; Figure S1). However most of these heterozygous introgressions were covered by another homozygous introgression in the same region in a different BIL. Nevertheless, although the BIL lines were not fully inbred after 4 or 5 generations of selfing, overall we found little residual heterozygosity (Figures 3A and 4). The comprehensive genotype of the 439 BILs along with the chromosome-wise distribution of introgressions for the population is presented in File S1 and File S2. In addition, the plots of the genotyped BILs are presented in an online resource for this paper (github.com/mfcovington/bil-paper).

**Figure 3.**
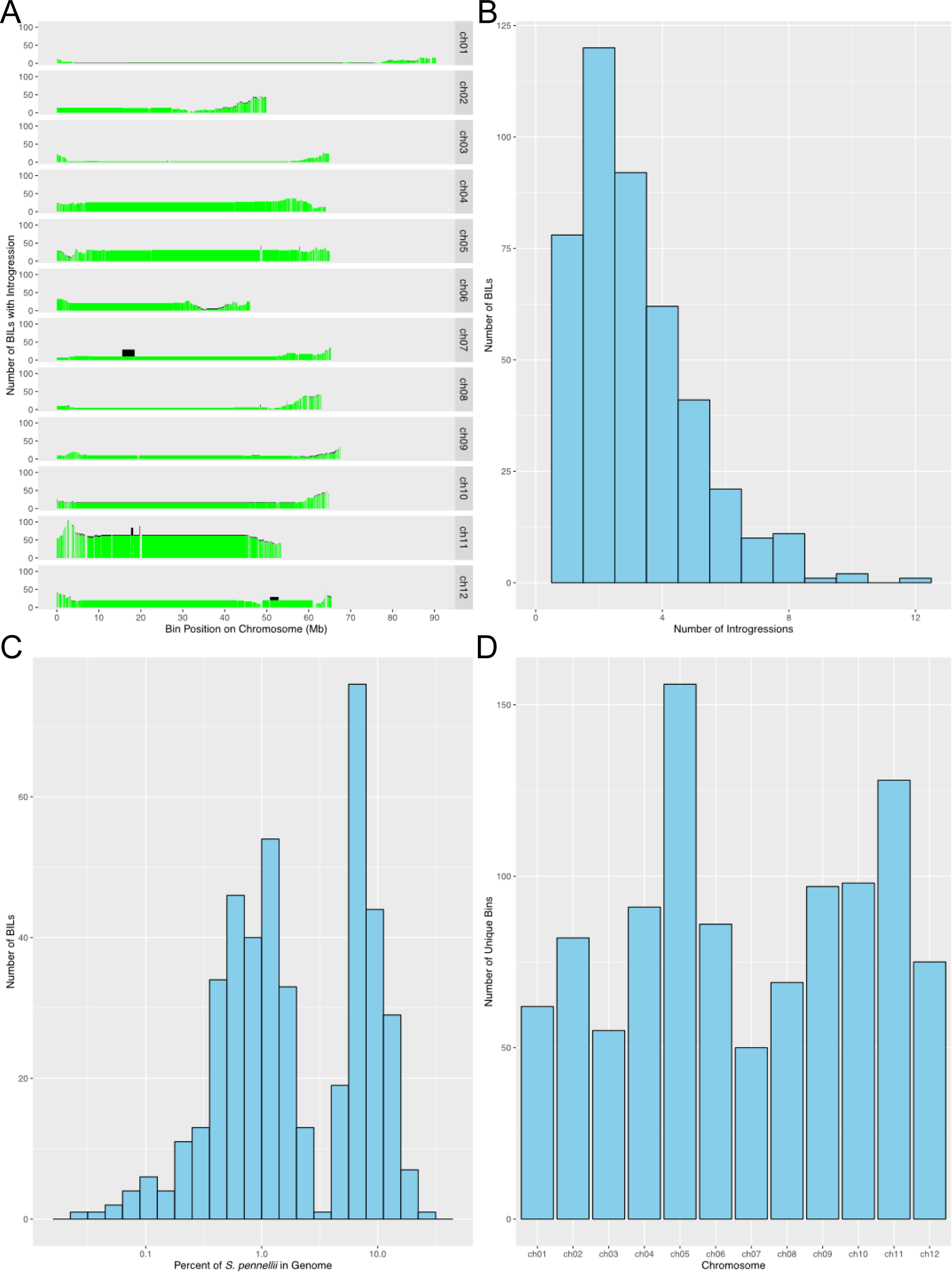
Introgression and bin statistics. A. Distribution of introgressions across bins. B. Introgression frequency per BIL. C. Introgression percentage per BIL. D. Number of bins per chromosome. Panel A is plotted in genetic distance as Supplemental Fig. S1.

The number of introgressions for each BIL varied from 1 to 12 with a mean of 3.125 introgressions per BIL (Figure 3B). Approximately 80% of the lines had from 1 to 4 introgressions (Figure 3B). As opposed to the single introgressions per line of the IL population, the presence of multiple introgressions in a single BIL enables the study of the genetic interactions regulating complex traits. The BIL population provides a synergistic complement to the IL population (Eshed and Zamir 1994), an additional resource for linking QTL identified in the ILs to genes as multiple BILs possess smaller overlapping introgressions encompassed within the larger introgression regions of the IL population. The mean and median numbers of genes per introgression are approximately 677 and 485 for the ILs, and 479 and 293 for the BILs. For example, the introgression in IL4-3 has been identified as having the largest contribution to tomato leaf traits in terms of the overall number of QTL mapped to this region (Holtan and Hake 2003; Chitwood *et al.* 2013). We identified at least 50 BILs with breakpoints overlapping the introgressed region of IL4-3, allowing fine-scale dissection of QTL in this region. We also explored the BIL population for the number of introgressions on specific chromosomes (Table 1). A relatively small number of BILs (54) have an introgression on chromosome 1, the largest chromosome by physical distance, whereas a surprisingly high number of BILs (236) have introgressions on chromosome 11 (Table 1; Figures 3 and 4). Chromosome 5 also harbors a large number of introgressions (171).

**Table 1.** Per chromosome BIL introgression and bin counts.

| **Chromosome** | **No. BILs with introgression** | **No. introgressions** | **No. bins** |
| --- | --- | --- | --- |
| **1** | 53 | 54 | 62 |
| **2** | 91 | 99 | 82 |
| **3** | 57 | 69 | 55 |
| **4** | 94 | 101 | 91 |
| **5** | 145 | 171 | 156 |
| **6** | 93 | 103 | 86 |
| **7** | 72 | 78 | 50 |
| **8** | 97 | 103 | 69 |
| **9** | 84 | 103 | 97 |
| **10** | 109 | 126 | 98 |
| **11** | 208 | 236 | 128 |
| **12** | 104 | 129 | 75 |

**Figure 4.**
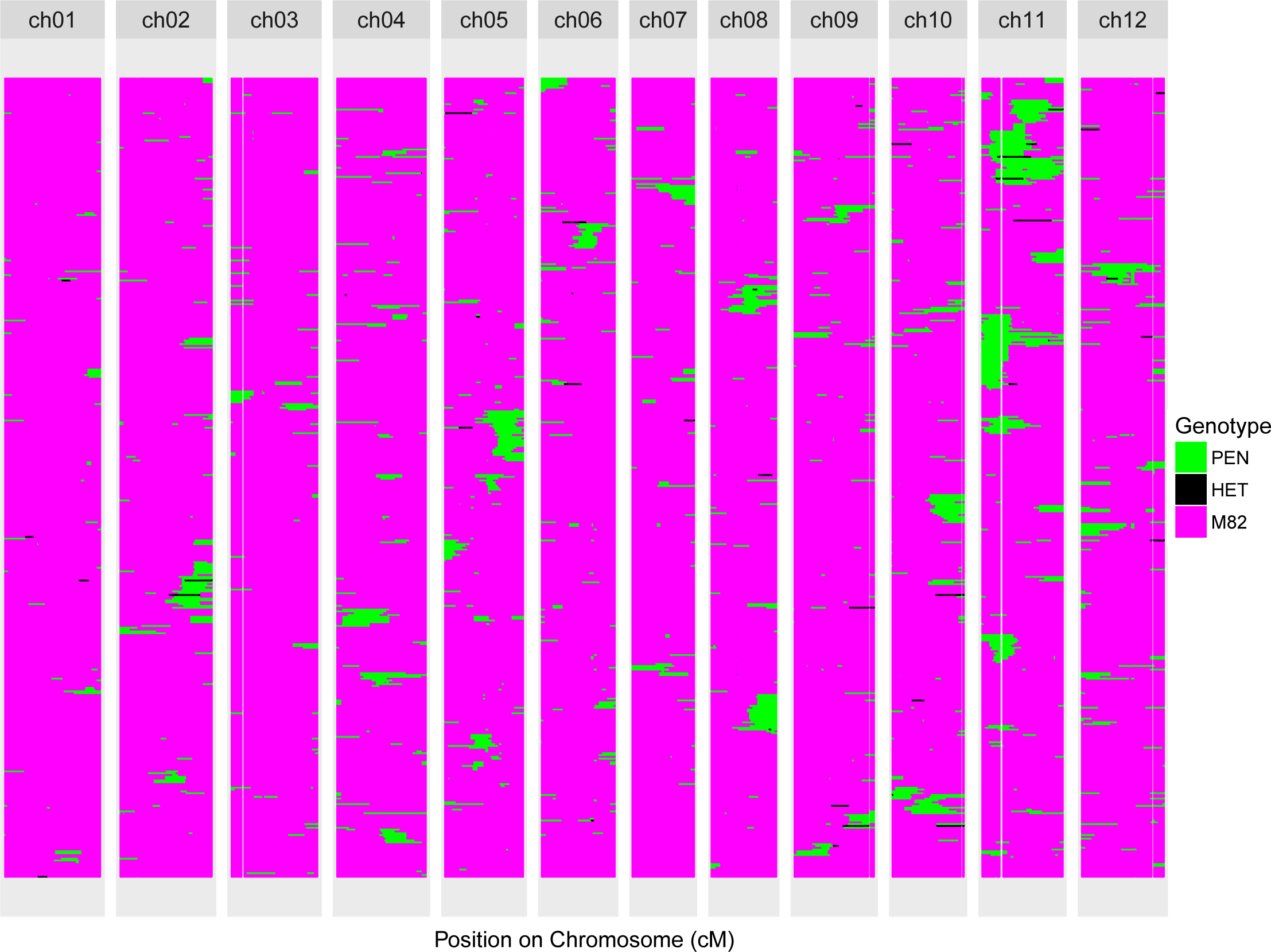
Composite genotype map of the entire BIL population. See Supplemental Fig. S2 for a version of this figure plotted by physical distance, and Supplemental Fig. S3 for a version plotted with BILs clustered by each chromosome after removing samples without an introgression.

The BILs showed remarkable variation in terms of the percentage of *S. pennellii* genome in each line (Figure 3C), ranging from <1% to >10%. The introgression size ranged from 16.7 Kb on chromosome 10 in BIL-025, containing only 4 genes, to the largest introgressions on chromosome 12 in BIL-083 and BIL-444, which span almost the entire chromosome and contain more than 2400 genes. Taken together, the remarkable variation in the number of introgressions, size of introgressions, and the level of genome contribution makes the BILs a highly useful resource for fine mapping of complex traits and identification of the underlying genes.

The precise boundary information for the genotyped BILs was used to define non-contiguous “bins”: regions defined by the pattern of overlap among introgressions in the population and within which no line changes genotype (Liu and Zamir; Chitwood *et al.* 2013). A total of 1049 unique bins (87 per chromosome on average) were detected across the entire genome with the maximum number of bins on chromosomes 5 and 11, with 156 and 128 bins respectively (Table 1; Figure 3D; File S3). Thus, the BILs provide an increase of nearly an order of magnitude in the number of bins as compared to the 112 bins defined in the IL population (Chitwood *et al.* 2013). We used our precision genotyping and introgression boundary calls to define the gene content of bins with near certainty. A list of annotated genes within the BIL bins is provided as File S4. Interestingly almost half of the bins defined for BILs (455 out of total 1049 bins) had 10 or fewer genes, which will greatly facilitate the identification of candidate genes in future studies. Conversely, only 9 out of 126 IL bins (at the sub-bin level) have 10 or fewer genes. For the ILs, the summary statistics for the number of genes per bin (at the sub-bin level) are: range = 1 - 1890, mean = 262.06, and median = 150. In contrast, for the BILs these values are: range = 1 - 984, mean = 23.15, and median = 7.

Although Ofner et al. (Ofner *et al.* 2016) used the same tomato BIL population, we achieved a finer resolution of recombination breakpoints and defined 66% more bins thanks to higher density genotyping (roughly 80 times greater number of SNPs). While merging the BIL introgression and bin maps presented in these two publications would be worthwhile, it is hampered by the fact that the population was genotyped at different stages in the two studies and that different versions of the tomato reference genome were used.

### Analysis of leaf trait variation in the population

As described above, the parents of this population show striking differences in leaf morphology and complexity. Not surprisingly, there is substantial variation for these same traits segregating in the BILs. In order to account for replication and pseudo-replication of trait measurements, linear mixed models (LMM) were used to estimate model-fitted BIL trait means for QTL mapping. As expected due to the back-crossed population design, the distribution of many of the traits is roughly centered around the recurrent parent, *S. lycopersicum* cv. M82, yet often present a skew towards the wild parent, *S. pennellii* (Figures S4-S29). This is most evident in traits such as flowering time and roundness, for which the only BILs significantly different from M82 are in the *S. pennellii* direction (Figures S13 and S11).

Overall, we found positive correlation among leaf complexity traits (i.e., the number of different types of leaflets and the total number of leaflets). There is stronger correlation between intercalary and secondary leaflet counts, than between either of these and primary complexity. Likewise, intercalary and secondary leaflets are also more strongly correlated than primary leaflets to the total number of leaflets (Figure S30). We found no correlation between leaf complexity and leaflet shape traits. Aspect ratio and roundness are very strongly negatively correlated, which is to be expected. Circularity and solidity are strongly correlated. In the context of tomato leaflet shape, dentation and lobing of the leaflet margin similarly affects circularity and solidity. As in our previous study of tomato leaflet shape in the ILs (Chitwood *et al.* 2013), we used an outline morphometrics technique, elliptical fourier descriptors decomposed into their principal components (EFD-PC), to examine leaflet shape in the BILs (Figures S14-S29). However, the EFD-PC traits showed low heritability and repeatability in the BILs, perhaps owing to the lower replication in this dataset (Table S1), so we excluded these traits from further discussion.

### QTL regulating leaf traits

We used the precisely defined BIL bins to identify QTL regulating various leaf developmental traits, such as complexity, aspect ratio, circularity, and solidity. Two methods, marginal regression (Chitwood *et al.* 2013) and SparseNet (Mazumder *et al.* 2011), were used to identify BINs associated with different traits. Hundreds of bins were significantly associated with complexity and leaf shape traits by marginal regression at a q-value of < 0.01 (File S5; Figures S31-S45). Corroborating what we found in the ILs, where ILs 4-3 and 5-4 harbored the largest number of bins associated with leaf traits (Chitwood *et al.* 2013), numerous BIL bins in chromosomes 4 and 5 were associated with leaf complexity and leaflet shape traits.

To increase mapping resolution beyond what is possible with marginal regression we used the regularized regression method SparseNet, which performs variable selection and shrinkage of the regression coefficients (i.e., inferred association between bins and traits). SparseNet reduced the number of trait associated bins by around an order of magnitude (Figure 5; Figures S46-S66; File S6). Only 28, 36, 26 and 7 bins were found to be associated with primary, intercalary, secondary, and total complexity, respectively. This method also identified a remarkably reduced number of bins and sharp QTL peaks for leaf shape traits. The bins associated with various leaf traits were distributed on multiple chromosomes across the tomato genome, emphasizing the polygenic regulation of leaf developmental traits.

**Figure 5.**
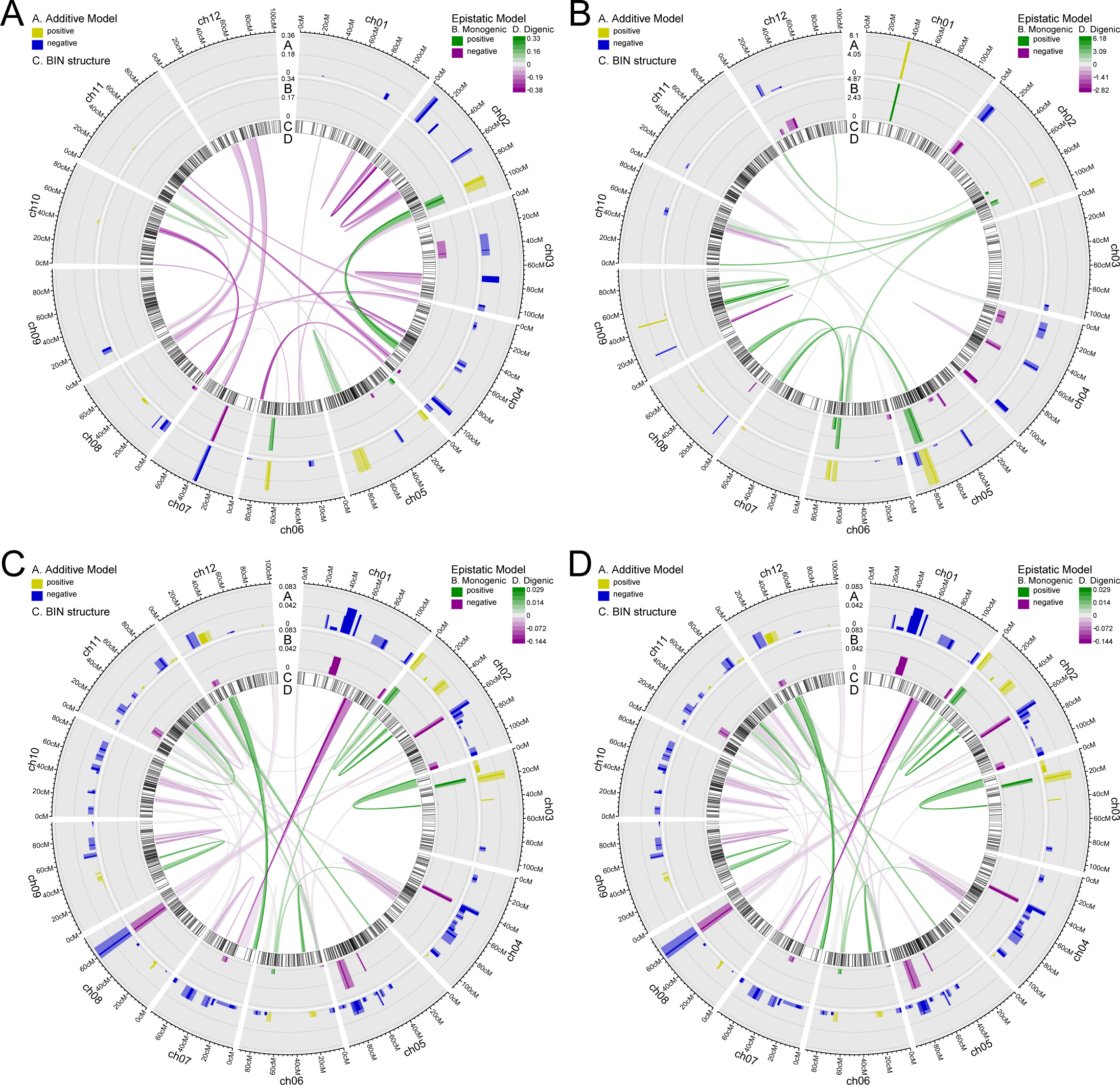
QTL mapping results for additive and epistatic models. A. Primary complexity.B. Total complexity.C. Aspect Ratio. D. Circularity. The bin selected by SparseNet as a QTL is colored opaquely, andthenearbybinscorrelatedtoitatorgreaterthan0.9aretakenasanapproximateQTL interval and colored translucently.

### Epistatic interactions for the leaf traits

Our previous work showed that tomato leaf complexity and leaflet shape is polygenic (Chitwood *et al.* 2013). This is also evident in the current study, where we detected numerous loci that contribute to leaf traits distributed over the entire tomato genome. Although functional genetic studies have reported evidence for epistasis in tomato leaf development (Koltai and Bird 2000; Ori *et al.* 2007; Naz *et al.* 2013), the genome-wide pattern of epistasis regulating tomato leaf traits has yet to be investigated.

The variable selection capability of SparseNet enabled us to search the full epistatic QTL model space without preconditioning on QTL with a main (i.e., single locus) effect. The 1049 BIL bins have 549,676 possible two-locus combinations, which define the set of potential epistatic QTL. We observed 306,291 (or roughly 56%) of said two-locus combinations in the BIL lines. Thus, the search space for the epistatic QTL model comprised 307,340 variables, 1049 single locus plus 306,291 two-locus variables. We inferred 25, 3, 33 and 23 epistatic interactions regulating primary, secondary, intercalary and total leaf complexity (Figure 5 A and B; Figures S46-S47; File S6). We further uncovered 42, 45, 53 and 61 possible epistatic interactions regulating aspect ratio, circularity, roundness and solidity of leaflets (Figure 5 C and D; Figures S48-S49; File S6). Although we did not do so, inclusion of a per-line effect in the regression model would help guard against false positive epistatic QTL in a backcrossed population such as the BILs with uneven number of lines covering different genomic regions. Similarly, higher replication per line would also lower the chance for false positives.

### Candidate genes underlying the detected QTL

In order to define candidate genes underlying the identified single locus and epistatic QTL, we used a previously established set of literature-curated leaf developmental genes (Ichihashi *et al.* 2014). We reasoned that any causal gene would have amino acid and/or *cis*-driven mRNA abundance differences segregating in the population. To identify genes with *cis*-driven mRNA abundance differences we filtered the literature-curated leaf developmental genes to retain those exhibiting *cis* eQTL using transcriptome profiling and expression-QTL analysis of the IL population (Chitwood *et al.* 2013; Ranjan *et al.* 2016). Out of 384 leaf developmental genes, 62 were detected with an underlying eQTL. Using these 62 genes, we identified roughly 10 candidate genes for each of the leaf complexity traits, and 10-20 genes for each leaf shape phenotype. *GROWTH-REGULATING FACTOR*, *ASYMMETRIC LEAVES 2*, *AINTEGUMENTA*, *ARGONAUTE*, *LEUNIG*, *LONELY GUY*, *STM*, *BEL1*, *TCP*, *YUCCA* and *IAA*were some of the salient candidate genes for both leaf complexity and shape traits (File S7). We used permutation analyses to determine the significance of enrichment of leaf developmental genes among the QTL identified. Significant enrichment of the literature-curated leaf developmental gene set was observed for leaf shape QTL, such as aspect ratio, circularity, solidity and roundness, but the enrichment was not significant for leaf complexity traits (Table S2). One possible explanation for this is that since the literature-curated leaf gene reference list was primarily based on Arabidopsis (simple) leaf development, it is likely biased for leaf shape and not representative of leaf complexity in tomato.

We also detected candidate genes underlying the epistatic QTL. One possible interacting pair of genes underlying an epistatic QTL for primary complexity is *GRF2* on chromosome 2 and *ANT* on chromosome 4 (Figure 5A; File S8). Surprisingly, our transcriptomic study of tomato, *S. pennellii*, and *S. habrochaites* compound leaf development found *GRF2* and *ANT* to be highly-connected hub genes at the core of the gene co-expression network (Ichihashi *et al.* 2014). That these independent analyses using different methodologies identified the same genes supports their importance for compound leaf development. This finding also underscores the importance of the regulation of cell proliferation for compound leaf development.

Another epistatic QTL for primary complexity with candidate genes for both loci was that between chromosomes 7 and 12, which contains *LSH10* and *TCP4* in one locus and *IAA CARBOXYMETHYLTRAFERASE1* in the other. *BEL1*, *ARGONAUTE 5* and *GRF4*were other strong candidate genes for epistatic QTL, but without obvious interacting candidates from the reference gene set on their paired interval. It should be noted that some of the epistatic QTL may in fact be broad-peaked single locus QTL, as the inferred pairs of interacting bins are on the same chromosome and possibly too close to have been separated by recombination in a few generations of crossing. Therefore, for the purpose of candidate gene identification for epistatic QTL we removed QTL whose bins are less than 20 cM apart. Following this filtering, we focused on the top 10 epistatic QTL by effect magnitude (File S8).

The above analysis only considered the candidate genes showing gene expression differences between the two parents and does not account for amino acid changes that could underlie the leaf shape and complexity differences between the two parents. Therefore, we used PROVEAN (Choi *et al.* 2012) to predict amino acid changes of functional significance between the two parents for the curated leaf developmental genes. This added 42 genes to the candidate genes reference list. Redoing the candidate gene search with an updated gene list, including both eQTL and PROVEAN data, increased the number of candidate genes for each trait but at the cost of significance in the permutation analysis (Table S2). Noteworthy PROVEAN candidate gene additions were *PIN-FORMED 1*, *GIBBERELLIN OXIDASE*s, *CLAVATA 1-like*, *IAA4*, *YUUCA8*, and *GRF5*.

Lastly, in order to explore if gene regulatory interactions underlie some of the inferred epistatic QTL we assessed whether IL *trans*-eQTL were enriched among the set of paired regions that make up the BIL epistatic QTL. We did this with two sets of *trans*-eQTL derived from eQTL clustering (Ranjan *et al.* 2016). The above set of literature-curated leaf development genes were highly represented in two eQTL clusters, one enriched for leaf development GO (gene ontology) terms and another enriched for photosynthesis GO terms. Therefore, we tested for IL *trans*-eQTL enrichment among BIL epistatic QTL by permutation using the leaf development cluster (set 1), as well as this cluster plus the photosynthesis cluster (set 2). Both sets yielded highly significant results with p-values of 0.0028 and < 1.0E-04, respectively for set 1 and 2 (Table S3).

### *Punctate:* a fine-mapping test case

To experimentally test the usefulness of the BILs in resolving genetic loci associated with a trait, we conducted a case study for the *Punctate* phenotype, which enhances the accumulation of anthocyanins in trichome bases creating a “purple spot” phenotype. Among ILs, IL10-3 shows the *Punctate*phenotype in leaves, a phenotype observed in *S. pennellii*but not M82. Notably, the Tomato Genetics Resource Center lists the *Punctate*locus as residing on chromosome 8 (see accessions LA0812, LA0998, and LA3089, tgrc.ucdavis.edu), suggesting a previously unknown locus on chromosome 10 is responsible for the trait variation segregating in the ILs and BILs. The *S. pennellii* introgression in IL10-3 spans 2.65 Mbp and 385 genes (Chitwood *et al.* 2013), making candidate gene identification difficult. To determine the extent to which the BILs could improve the mapping resolution, we screened the BILs and identified 54 lines with the phenotype. Of the 54 BILs exhibiting the “purple spot” phenotype, 46 possessed an introgression on chromosome 10 that overlapped the IL10-3 introgression. The remaining 8 BILs did not show the *Punctate* phenotype consistently. Among BILs with a chromosome 10 introgression and the “purple spot” phenotype BIL-228 had the smallest introgression, covering 310 Kbp and containing only 51 genes, reducing the number of candidate genes from 385. By examining BILs with chromosome 10 introgressions yet without the *Punctate* phenotype, we further narrowed the associated genomic region to a 114 Kbp segment with only 18 genes (Figure 6). Thus, the BILs enabled us to dramatically reduce the region and the number of candidate genes for the “purple spot” phenotype by 96% and 95% respectively.

**Figure 6.**
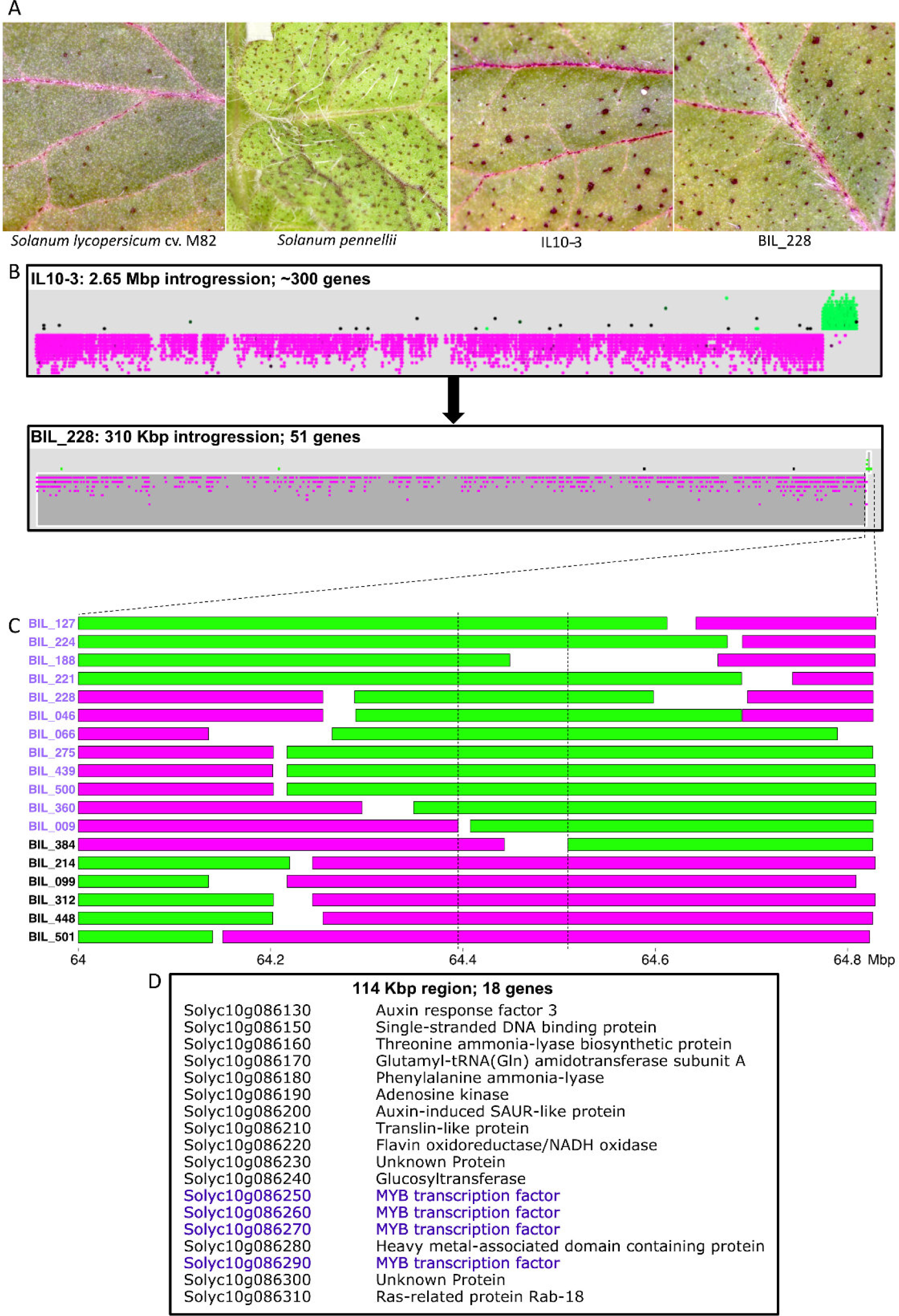
Punctate fine-mapping. A. Adaxial leaf images showing presence and absence of the “purple spot” phenotype. B. Comparison of the chromosome 10 introgressions of IL10-3 and BIL-228. C. Fine-mapping the *Punctate* locus using BILs; purple-labeled BILs possess the “purple spot” phenotype, whereasblack-labeledBILsdonot.D.Genelistofthefine-mappedregionwithcandidate MYB tandem expansion highlighted in purple.

Among the 18 genes in the *Punctate* fine-mapped region, there were four tandemly-duplicated genes encoding MYB transcription factors. Overexpression of an Arabidopsis homolog of these MYB genes (MYB114, AT1G66380), part of an independently-derived tandem duplication of MYB genes that regulates anthocyanin biosynthesis, results in a *Punctate*-like phenotype with accumulation of anthocyanins in trichome bases (Gonzalez *et al.* 2008). Remarkably, a similar tandem duplication of four MYB homologs in grape (*Vitis vinifera*) (*VvMybA1, VvMybA2, VvMybA3*) explains 62% of the variation of anthocyanin accumulation in berries in a Syrah x Grenache F1 QTL study (Fournier-Level *et al.* 2009). This reinforces the involvement of MYB transcription factors in the regulation of the *Punctate* phenotype in *S. pennellii* leaves, and the importance of tandemly duplicated homologs found in lineages as diverse as *Arabidopsis*and *Vitis vinifera* in regulating anthocyanin accumulation in a tissue-specific manner. Similarly, BILs could facilitate the identification and cloning of genes regulating various other developmental and metabolic traits.

### Flowering time QTL and candidate genes

We also used the BIL population to infer QTL for flowering time and identify their candidate genes. Given the well-characterized pathways and genes for flowering time, this provides a good opportunity to further test the utility of the BIL population for candidate identification. Marginal regression identified 69 significant bins associated with flowering time in tomato. Interestingly, 49 of those were contiguous bins on chromosome 5 (Figure S39). The remaining bins were localized on chromosomes 8, 9 and 12. Three narrow QTL, one each on chromosomes 5, 8 and 12, were found to be associated with flowering time using the more stringent SparseNet method (Figure 6B, ring A). *SELF-PRUNING 5G (SP5G)*, a *FLOWERING LOCUS T* homolog, is present in the QTL on chromosome 5 and is the strongest candidate gene, by virtue of its function, that we identified for flowering time in the BILs (Carmel-Goren *et al.* 2003). The QTL on chromosome 12 also has some important developmental genes within it, such as *AP2*, *ULT* and *YABBY*, that are plausible candidates for the regulation of flowering time in tomato.

The QTL on chromosome 5 is the major flowering time QTL in the BILs by effect size and is coincident with a large effect QTL in the ILs (Figure 7A, ring C). The overlap between the BIL and IL QTL on chromosome 5 defines a 297-Kbp region containing 37 genes, among which *SP5G* is the only flowering time gene (Figure 6B, inset E; Table S4; Jones *et al.* 2007). Thus, the *SP5G* example not only highlights the utility of the BILs for fine-mapping, but also the advantage of using the BILs in conjunction with the ILs for QTL confirmation and fine-mapping.

**Figure 7.**
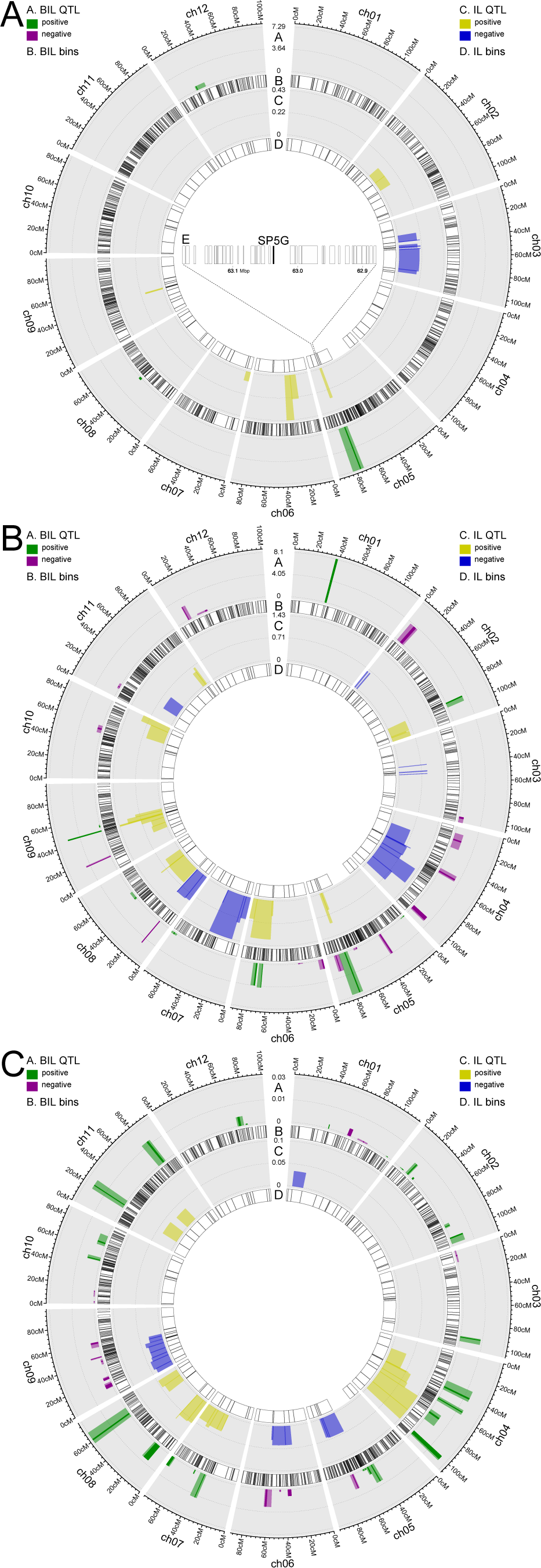
Fine-mapping and confirmation of IL QTL using BILs. A. Flowering time, showing the candidate gene *SP5G* in inset E. B. Total complexity.C.Circularity. The BIL bin selected by SparseNet as a QTL is colored opaquely, andthenearbybinscorrelatedtoitatorgreaterthan0.9 are taken as an approximate QTL interval and colored translucently.

### QTL confirmation and fine-mapping

Beyond the above two test cases, we explored the benefit of combining results from the IL and BIL populations for the purposes of QTL confirmation and fine-mapping since we measured the same traits in both populations. As mentioned above, the IL and BIL populations share the same parents, but have different crossing schemes and broadly disparate numbers of lines. The ILs have a near isogenic line (NIL) design, whereas the BILs have a mixed *BC*_2_ / *BC*_3_ advanced backcross design.

One advantage of utilizing both of these populations is that QTL can be validated in an independent experiment using a different set of lines. In our case, we found broad congruence between the IL and BIL QTL mapping results (Figure 7; Figures S67-S72; File S9). Nevertheless, there are discrepancies in the QTL inferred from these two populations, which could be ascribed to their different crossing schemes and patterns of wild introgression, statistical power differences due to disparate sample sizes, and false positives. Apart from the *Punctate* example several IL QTL were significantly narrowed in the BILs, such as QTL for total leaf complexity on chromosomes 8 and 9 (Figure 7B), circularity QTL on chromosomes 8 and 11 (Figure 7C), and several QTL for primary, secondary, and intercalary complexity (Figures S67-S69). Likewise, several recent studies have used a subset of the BILs with introgressions in a region of interest to further fine-map a QTL found in the ILs (Ning *et al.* 2015; Müller *et al.* 2016; Fan *et al.* 2016).

Furthermore, the use of these two populations is quite complementary. The ILs allow for higher replication of trait measurements due to the lower number of lines. NIL populations, such as the ILs, have also been shown to enable the detection of smaller effect QTL (Keurentjes *et al.* 2007). In contrast, the BILs can greatly facilitate higher resolution mapping (i.e., narrower QTL intervals) due to their generally narrower bins. Moreover, the BILs enable the mapping of epistatic QTL, which is not possible with the ILs. Our ability to infer epistatic QTL was further empowered by our high precision genotyping HMM and use of the regularized regression method SparseNet.

### Conclusions

The BILs provide a valuable new resource for the tomato research community, one that is complementary to the long-established IL population yet also extends the research toolkit considerably. We demonstrated the utility of this new population for corroborating and fine-mapping IL QTL. Furthermore, we established the capacity of the BIL population for epistatic QTL mapping and for revealing the genetic architectures of tomato traits. The present study also highlights the dramatic gains that can be reaped from the use of advanced statistical techniques, such as our novel heterogeneous genotyping HMM and the application of regularized regression for QTL mapping.

## Supplemental datasets and weblinks

**Weblink S1. Recombination boundary plots:** github.com/mfcovington/bil-paper/data/plots-with-boundaries/

**Weblink S2. Recombination boundary files:** github.com/mfcovington/bil-paper/data/boundaries/

**File S1. Introgressions by BIL.** Number and location of introgressions organized by BILs.

**File S2. Introgressions by chromosome.** Number and location of introgressions organized by chromosome.

**File S3: Bin locations and genotypes.** Bin locations and genotypes of the BILs at each bin ordered by chromosome.

**File S4: Annotated genes per BIN.** Genes organized by bins and with functional annotations.

**File S5: Marginal regression QTL mapping results**. There is one table per trait. QTL are defined as bins significantly correlated to traits by marginal regression at a BH-adjusted p-value of less than 0.01.

**File S6: SparseNet QTL mapping results.** There is one table per trait, empty if there were no QTL found for a given trait. The SparseNet selected bin is shown for each QTL, along with its coordinates, the estimated regression coefficient, and the 0.95 and 0.9 bin correlation intervals. For the epistatic QTL this information there are two selected bins and two sets of correlated intervals.

**File S7: Single locus QTL candidate genes.** There is one table per trait. The SparseNet selected bin is shown, the ITAG gene number for the candidate genes, the gene coordinates, whether the gene has PROVEAN or *cis*-eQTL signal, and the closest Arabidopsis homolog of the candidate gene.

**File S8: Epistatic QTL candidate genes.** There is one table per trait. The SparseNet selected bins are shown, together with their 0.9 bin correlation intervals, and other data from the mapping results. For QTL with candidate genes, subsequent lines in the tables show: the bin that contains the candidate gene, its ITAG gene number, chromosome, gene coordinates, expression difference in the ILs with respect to M82, the type of eQTL (*trans* vs *cis*), and the closest Arabidopsis homolog of the candidate gene.

**File S9: IL-BIL fine-mapping tables.** There is one table per trait showing overlapping QTL between the IL and BIL results. For the IL QTL the bin and sub-bin are shown, the bin coordinates, chromosome, slope (i.e., effect size), and significance values. For the BIL QTL the SparseNet selected bin is shown for each QTL, along with its coordinates, the estimated regression coefficient, and the 0.95 and 0.9 bin correlation intervals. For the epistatic QTL this information there are two selected bins and two sets of correlated intervals.

## Acknowledgments

We thank Yaniv Brandvain and Paul Lott for discussions and assistance with early versions of the HMM. We thank the Vincent J. Coates Genomics Sequencing Laboratory at UC Berkeley (supported by NIH S10 Instrumentation Grants S10RR029668 and S10RR027303), and computational resources/cyber infrastructure provided by the iPlant Collaborative (www.iplantcollaborative.org), funded by the National Science Foundation (Grant DBI-0735191). This work was supported through a National Science Foundation grant (IOS-0820854) awarded to NRS and JNM. DHC was a fellow of the Life Sciences Research Foundation funded through the Gordon and Betty Moore Foundation.

## Author’s contributions

DHC, JNM and NRS conceived and designed the experiments. DF, AR, DHC, DW, YS and LH performed the experiments. DF, AR, MFC, and DHC analyzed the data. DF developed new statistical inference tools. DF, AR, IF, MFC, DHC, DZ, JNM contributed reagent/materials/analysis tools. DF, AR, MFC, DHC, JNM, NRS wrote the paper. All authors read and approved the final manuscript.

